# Transcriptomic Analysis of CAD Cell Differentiation

**DOI:** 10.1101/2025.03.09.642086

**Authors:** Carlos A. Cevallos, Anna Leigh White, Brooke A. Fazio, Lillian S. Wendt, Jasmine W. Feng, Dora Posfai, April L. Horton, John M. Warrick, Omar A. Quintero-Carmona

**Author notes:** Authors contributed equally (Carlos A. Cevallos, Anna Leigh White, Brooke A. Fazio).

## Abstract

CAD cells were derived from Cath.a cells, a mouse central nervous system catecholaminergic cell line. Serum-starved CAD cells undergo morphological changes and resemble isolated neurons when observed by microscopy. We carried out an RNAseq transcriptomic analysis to examine differentiated CAD cells for expression signatures related to neuronal functions, identifying ∼1900 transcripts whose expression changed with differentiation. Pathview analysis identified ∼80 KEGG pathway gene sets that were differentially expressed, including upregulation of at least 13 neuron-related pathways. This dataset can be explored more deeply, allowing further investigation into expression changes relevant to studying neuronal functions in this easy-to-culture model system.

## Introduction

Neuron differentiation is difficult to study with primary neuronal cultures due to their delicacy and dependence on supporting glial cells. Most immortalized nerve cell lines also present challenges as proliferating neurons do not differentiate, making them unsuitable as neuronal models. <u>C</u>ath.<u>a</u>-<u>d</u>ifferentiated (CAD) cells, derived from the CNS catecholaminergic cell line Cath.a (Suri et al., 1993), present a possible solution to this issue (Qi et al., 1997). While CAD cells proliferate in serum-containing media, they differentiate, growing neurite-like processes when switched to serum-free media. CAD cells can be maintained for at least 6 weeks in differentiating media, making them an attractive model for studying neuronal processes and differentiation, especially for researchers where primary neurons are not easily accessible, as is the case for researchers at primarily undergraduate institutions.

The Cath.a cell line is derived from a transgenic mouse brain tumor carrying the SV40 T antigen (Tag) oncogene (Suri et al., 1993). Tag stimulates cell proliferation and likely blocks differentiation by keeping Cath.a cells in cycle (Fanning and Knippers, 1992). Like Cath.a, CAD cells express tyrosine hydroxylase (*Th*) and are generally accepted to be catecholaminergic. Unlike Cath.a, the CAD cell line produces no dopamine or norepinephrine despite expressing *Th* and instead exhibits large buildups of L-dihydroxyphenylalanine (L-DOPA), the product of TH enzymatic activity on tyrosine and the precursor to dopamine and norepinephrine, suggesting issues with DOPA decarboxylase activity. Additionally, CAD cells have lost the Tag gene and have gained the ability to differentiate under appropriate conditions (Qi et al., 1997).

Initial characterization by Qi and colleagues demonstrated that CAD expresses neuron-specific proteins such as class III β-tubulin, an isotype of tubulin expressed only in neurons (Sullivan et al., 1986), GAP-43, a cortically associated protein found at high concentration in axons and growth cones (Skene et al., 1986), SNAP-25, a presynaptic protein necessary for vesicle endocytosis (Oyler et al., 1991), and synaptotagmin, a synaptic vesicle protein (Matthew et al., 1981). Additionally, Qi et al. demonstrated that CAD cells do not express glial fibrillary acidic protein (GFAP), indicative of its neuron-like nature as many immortalized neuron lines express both glial and neuronal proteins (Schubert et al., 1974). Structurally, Qi et al. identified that CAD cells processes contain many of the same elements as neurons, including cytoskeletal elements, the presence of mitochondria, the absence of other organelles, dense-core and clear vesicles, and varicosities. Previous works have looked at changes in expression for subsets of genes related to neuronal functions (Bisig et al., 2009; Lazaroff et al., 1996; Suri et al., 1993), but a comprehensive assessment of gene expression in response to differentiation has not existed for CAD cells.

## Results and Discussion

To better understand the neuron-like nature of differentiated CAD cells, we conducted global examination of changes in gene expression resulting from CAD cell differentiation. In addition to examining gene expression changes for individual genes, we also investigated expression changes in gene systems as described by the <u>K</u>yoto <u>E</u>ncyclopedia of <u>G</u>enes and <u>G</u>enomes (KEGG) (Kanehisa and Goto, 2000). We differentiated CAD cells by growing them in serum-free media, and after 5 days the CAD cells produced long projections that ultrastructurally resembled neurites when visualized by fluorescence microscopy (Figure 1A) or scanning electron microscopy (Figure 1B). These processes contained mitochondria and terminated in structures resembling actin-rich growth cones.

**Figure 1:**
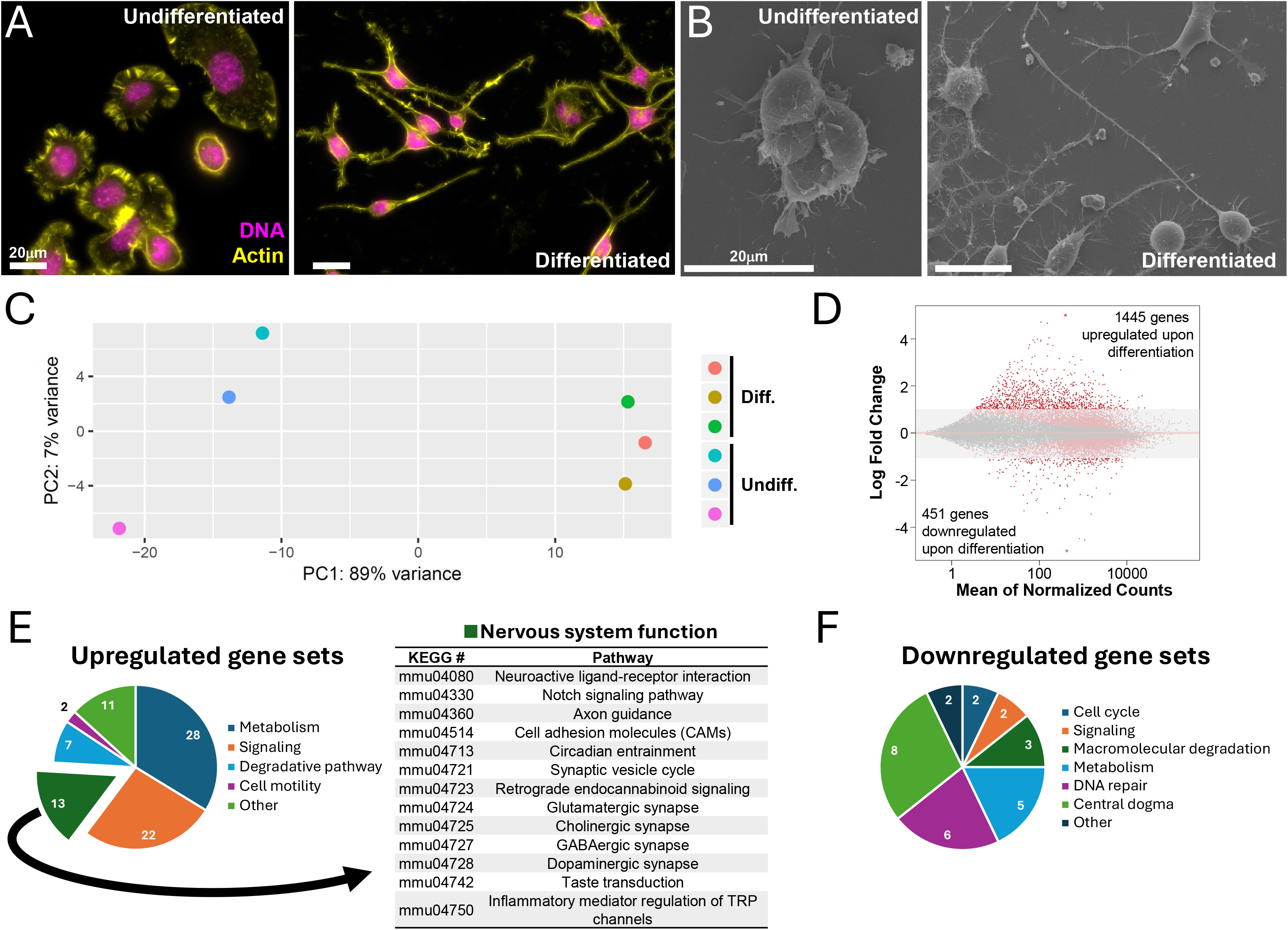
CAD cells differentiate into neuron-like cells with serum deprivation. The cell morphology of CAD cells changes to become more neuron-like after 5 days of growth in serum-free media. Undifferentiated cells form lamellae and sit flat or in clumps, while differentiated cells have long projections that form networks. (A) Phalloidin staining for actin is shown in yellow and DAPI staining for DNA is shown in magenta as visualized by fluorescence microscopy. (B) Representative scanning electron microscopy images. Displayed images are representative of commonly observed phenotypes. All scale bars are 20 μm. (C) Principal component analysis (PCA) indicated that 89% of the variation between RNAseq samples of differentiated and undifferentiated CAD cells could be attributed to differentiation state of the cells. (D) After 5 days of differentiation, 1445 genes were upregulated and 451 genes were downregulated with a more than 2-fold change in gene expression. Red markers outside of the shaded area indicate expression-level changes greater than 2-fold in magnitude with a p_adj_ ≤ 0.1. (E) Pathview analysis revealed differential expression for a subset of KEGG pathway gene sets compared to undifferentiated CAD cells, including upregulation of 13 gene pathways related to neuronal function (q ≤ 0.1).

We examined gene expression patterns of differentiated and undifferentiated cells by isolating RNA from cultures of cells and then completing an RNAseq analysis. Principal component analysis (PCA) revealed robust biological replication with undifferentiated and differentiated samples clustering together. The PCA also suggested up to 89% of variation between the transcriptome of samples can be attributed to the differentiated or undifferentiated state of the cells (Figure 1C). Upon differential gene analysis, 1445 genes were found to be significantly upregulated > 2-fold and 451 downregulated > 2-fold in differentiated cells compared to undifferentiated cells (p_adj_<0.1). Of the 1445 upregulated genes, 1050 had a baseMean value greater than 20. The baseMean values for *Actb* (100,911), *GAP34* (7203), *Tubb3* (16,487), *Snap25* (5,365), and *Myo10* (267) demonstrate a wide range of expression levels for genes relevant to neuronal function. The same was observed within gene families, such as the synaptotagmins, where baseMean values ranged from 30 (*Synt9*) to ∼7000 (*Synt1*). The gene expression data are included in the extended data. The full dataset discussed in this publication have been deposited in NCBI’s Gene Expression Omnibus (Edgar et al., 2002)and are accessible through GEO Series accession number GSE291553 (https://www.ncbi.nlm.nih.gov/geo/query/acc.cgi?acc=GSE291553).

RNAseq analysis revealed that many of the upregulated genes are implicated in all aspects of neuronal activities. For example, action potential firing has been observed in CAD cells (Wang and Oxford, 2000), and differentiated CAD cells displayed increased expression of a subset of the subunits for voltage-gated sodium channels such as *Scn1b* (baseMean=46, 6.3-fold increase, p_adj_<1.1×10^-11^) or *Scn3a* (baseMean=1344, 4.3-fold increase, p_adj_<1.2×10^-31^), potassium channels such as *Kcnab1* (baseMean=1838, 2-fold increase, p_adj_<4.5×10^-20^) or *Kcnc1* (baseMean=923, 2.8-fold increase, p_adj_<0.001), calcium channels such as *Cacna1b* (baseMean=1590, 2.2-fold increase, p_adj_<3.3×10^-5^) or *Cacna2d2* (baseMean=571, 2.7-fold increase, p_adj_<0.012), and sodium/potassium pumps such as *Atp1b2* (baseMean=491, 2.7-fold increase, p_adj_<5.6×10^-10^) (Bean, 2007; Bezanilla, 2008; Burke and Bender, 2019; Rama et al., 2018).

Expression of tyrosine hydroxylase (*Th*) does not change significantly upon differentiation (baseMean=91, 1.3-fold expression change, p_adj<_0.3). Although, dopa decarboxylase (*Ddc*) expression increases approximately 2-fold (baseMean=60, 2-fold expression increase, p_adj_<0.03) upon differentiation, previous data indicate that this change is insufficient to result in appreciable levels of dopamine expression (Qi et al., 1997). This could be due to posttranscriptional contexts that limit dopamine synthesis, including translational regulation, cellular conditions insufficient for the DDC enzyme to be sufficiently active for dopamine synthesis. *Ddc* is also the second enzyme in the serotonin synthesis pathway, while the first enzyme in the pathway, tryptophan hydroxylase, showed a decrease in expression (*Tph2*, baseMean=146, ∼40% decrease in expression, p_adj_<4.8×10^-4^). Following neurotransmitter synthesis, these small molecules are loaded into synaptic vesicles by membrane transporters such as *Slc17a7/Vglut1* (baseMean=47, 3.8-fold expression increase, p_adj_<2.4×10^-7^). This glutamate transporter is found in synaptic vesicle membranes (Sanchez-Mendoza et al., 2017), is one of the most highly expressed glutamate transporters in the adult brain (Wojcik et al., 2004) and is preferentially expressed in neuroblasts over glial cells (Sanchez-Mendoza et al., 2017).

Differentiation also resulted in a significant increase in receptors involved in synaptic function, such as dopamine receptor D1 (*Drd1*, baseMean=55, 4-fold increase, p_adj_<3.7×10^-8^), in agreement with previous observations (Pasuit et al., 2004). Proteins involved in synapse establishment (Craig and Kang, 2007) were also upregulated, such as the pre-synaptic cell adhesion molecule neurexin 1 (*Nrxn1*, baseMean=289, 2.5-fold increase, p_adj_<1.8×10^-11^) the post-synaptic protein neuroligin 2 (*Nlgn2*, baseMean=1848, 1.8-fold increase, p_adj_<1.6×10^-4^). Neurotransmitter reuptake attenuates neuronal signaling. *Slc6a17*, is a glutamate transporter present at synaptic junctions in both pre- and post-synaptic sites, particularly in glutamatergic terminals (Iqbal et al., 2015), and was upregulated approximately 8-fold in in differentiated CAD cells (baseMean=18, p_adj_<3.4×10^-6^).

In addition to these specific genes involved in neuronal activity, Pathview analysis (Luo and Brouwer, 2013) revealed upregulation for entire pathways as well. Gene enrichment analysis and functional classification by KEGG revealed 83 gene sets that are significantly upregulated and 28 that are downregulated in differentiated cells (q*<*0.1). The pattern of upregulation for neurotransmitter biosynthetic enzymes, neurotransmitter receptors, and other related machinery varied across the different pathways. The full Pathview analysis is included in the extended data.

Thirteen of the upregulated gene sets were related to neuronal function (Figure 1F). Genes in the glutaminergic (mmu04724, q=4.0×10^-2^), cholinergic (mmu04725, q=4.9×10^-5^), GABAergic (mmu04727, q=1.6×10^-3^), and dopaminergic (mmu04728, q=2.1×10^-2^) synapse pathways were all upregulated, as well as the synaptic vesicle cycle pathway (mmu04721, q=1.1×10^-2^), suggesting that differentiated CAD cells possess a subset of the machinery required for some forms of neuronal communication. Other KEGG pathways related to extracellular signaling relevant to neuronal function were also upregulated, including neuroactive ligand-receptor interaction (mmu0480, q=2.3×10^-2^), retrograde endocannabinoid signaling (mmu04723, q=5.1×10^-2^), and Notch signaling pathway (mu04330, q=3.3×10^-2^). Additional upregulated pathways included systems related to migration, such as cell adhesion molecules (mmu04514, q=4.9×10^-5^), ECM-receptor interaction (mmu04512, q=1.4×10^-4^), focal adhesion (mmu04510, q=3.1×10^-3^) and axon guidance (mmu4360, q=2.3×10^-3^).

Pathview analysis also revealed the downregulation of pathways less prevalent in mature neurons, including DNA replication (mmu03030, q=2.1×10^-25^) and cell cycle (mmu04110, q=1.5×10^-23^). Additionally downregulated were several pathways related to DNA repair including homologous recombination (mmu3440, q=6.2×10^-14^), mismatch repair (mmu03430, q=2.7×10^-9^), nucleotide excision repair (mmu03420, 4.0×10^-9^), and non-homologous end-joining (mmu03450, q=4.7×10^-2^). These findings align with Qi et al.’s observation that initiating differentiation slowed CAD cell proliferation (Qi et al., 1997). Taken together, the downregulation of these pathways is indicative of a population of cells no longer actively dividing, as would be the case for neurons and other terminally differentiated cells (Anda et al., 2016). However, transcriptionally active, long-lived cells such as neurons require repair mechanisms to maintain genomic integrity (Li et al., 2022), so the downregulation of nucleotide excision repair (mmu03420, q=4.0×10^-9^) and base excision repair (mmu03410, q=1.3×10^-5^) pathways is surprising.

A variety of model systems exist for studying neuronal function at the scale of individual cells, such as isolated primary neurons (Jiang et al., 2006; Mains and Patterson, 1973), induced pluripotent stem cells (Compagnucci et al., 2014; Dolmetsch and Geschwind, 2011; Inoue, 2010; Swistowski et al., 2010), and a variety of cultured cell lines (Biedler et al., 1973; Jones-Villeneuve et al., 1982; Pleasure et al., 1992; Schubert et al., 1974; Schubert et al., 1977). Many of these systems are expensive to maintain and challenging to grow in culture. Additionally, some of the cell lines are not derived purely from the CNS (Wang and Oxford, 2000). The CAD cell transcriptome suggests that there are gene-specific, pathway-specific and global similarities in gene expression between differentiated CAD cells and mature neurons. In conclusion, CAD cells provide an interesting and viable option for researchers to employ a relatively easy-to-culture model system with many of the structural features of neurons. This transcriptomic dataset allows researchers to determine if their particular research question could be investigated using CAD cells, based on examination of expression changes in relevant genes and pathways.

## Acknowledgements

We thank the Duke University School of Medicine for the use of the Sequencing and Genomic Technologies Shared Resource, which provided library generation, RNA sequencing services, and the guidance to make initiation of this project possible. Thank you to the other members of the Q-lab for their enthusiasm and support in pursuit of new knowledge. The University of Richmond School of Arts & Sciences funded undergraduate summer research for Anna Leigh White, Brooke Fazio, Lillie Wendt, and Jasmine Feng through “The Richmond Guarantee.” This work was also funded by NIH: R15GM119077 to Omar Quintero-Carmona and R15GM119077-S1 (Research Supplements to Promote Diversity in Health-Related Research) which supported Carlos Cevallos’s work as a post-baccalaureate researcher.

## Author contributions

Carlos A. Cevallos: Funding acquisition, Formal analysis, Investigation, Visualization, Writing – original draft

Anna Leigh White: Formal analysis, Investigation, Visualization, Writing – original draft, Writing – review & editing

Brooke A. Fazio: Formal analysis, Investigation, Writing – original draft

Lillian S. Wendt: Visualization, Writing – original draft, Writing – review & editing

Jasmine W. Feng: Formal analysis, Visualization, Writing – original draft, Writing – review & editing

Dora Posfai: Conceptualization, Data curation, Formal analysis, Investigation, Methodology, Resources, Software, Validation, Visualization, Writing – original draft, Writing – review & editing

April L. Horton: Funding acquisition, Conceptualization, Supervision, Writing – original draft, Writing – review & editing

John M. Warrick: Supervision, Writing – original draft, Writing – review & editing

Omar A. Quintero-Carmona: Conceptualization, Data curation, Formal analysis, Funding acquisition, Investigation, Methodology, Project administration, Resources, Supervision, Validation, Visualization, Writing – original draft, Writing – review & editing

## Materials and Methods

### CAD cell culturing

Cath.a-differentiated (CAD) cells were a gift from Dona Chikaraishi (Duke University) (Qi et al., 1997), and were maintained in growth media (DMEM/F12 media supplemented with 10% fetal bovine serum, penicillin, and streptomycin) in a humidified atmosphere containing 5% CO_2_ at 37°C. To induce differentiation, cells were transitioned to differentiating media (DMEM/F12 media supplemented with penicillin, streptomycin, and ITS supplement (Gemini) containing insulin, transferrin, and selenium).

### Scanning electron microscopy

Acid-washed coverslips were coated with 10μg/ml mouse laminin in PBS for 1 hour and washed with PBS to remove unbound laminin. Cells were plated on coverslips in growth media at a density appropriate for imaging individual cells and allowed to adhere overnight. Samples were fixed in PBS with 2.5% glutaraldehyde for a minimum of 20 minutes. The coverslips were then washed in PBS followed by a series of washes in increasing concentrations of ethanol for 5 minutes at each step (30%, 50%, 70%, 90%, and 100%). The samples were then critical-point dried and sputter-coated as described previously (Applewhite et al., 2016). Cells were imaged using a JEOL 6360LV scanning electron microscope. For differentiated samples, cells were transitioned to differentiating media for five days prior to processing for imaging.

### Fluorescence microscopy and immunostaining sample preparation

Cells were plated on acid-washed, 22mm square #1.5 coverslips which had been previously coated with 10μg/ml mouse laminin in PBS for 1 hour. Coverslips were seeded at a cell density appropriate for imaging and allowed to adhere overnight in growth media. Differentiated samples were transitioned to differentiating media for five days prior to processing for imaging. For simple staining for DNA, and F-actin, samples were fixed in 37°C PBS containing 4% paraformaldehyde for 10 minutes, and then permeabilized in PBS containing 0.5% Triton X-100 for 5 minutes. Samples were then stained in PBS containing 6.6nM ALEXA488 phalloidin and 100nM DAPI for 20 minutes. The coverslips were then washed 4 times in PBS for 5 minutes and mounted on slides in PBS containing 80% glycerol and 0.5% N-propyl gallate, and sealed with clear Sally Hansen Xtreme Wear nail polish.

### Fluorescence microscopy image acquisition

Images were acquired using an Olympus IX-83 microscope outfitted with a PLAN APON 60x/1.42NA DIC objective, or a UPlanFL N 40x/1.3NA DIC objective. Fluorescence imaging required an EXFO mixed gas light source, Sutter filter wheels and shutters, and a Sedat Quad filterset (Chroma). Images were acquired using a Hamamatsu ORCA-Flash 4.0 V2 sCMOS camera, and Metamorph imaging software. Images were prepared for publication using Metamorph and FIJI/ImageJ (Schindelin et al., 2012), which included brightness adjustment, contrast adjustment, and pseudocoloration.

### RNA isolation

RNA was isolated from T25-sized cultures of CAD cells using the RNeasy Plus kit (Qiagen) according the supplied protocol for purification of RNA from animal cells, with the addition of the Qiashredder step prior to loading the lysate on the column as well as the addition of an on-column DNAse digestion prior to washing and eluting the RNA from the column. RNA was then frozen and sent to the Duke Center for Genomic and Computational Biology for further processing and sequencing.

### Library-generation and RNA sequencing

The Duke Sequencing and Genome Technologies Core built stranded mRNA-seq libraries from RNA using Kapa Stranded RNA HyperPrep Kits (Roche). Libraries were pooled in equimolar concentration and were sequenced on the Illumina HiSeq 4000 producing 50 bp single-end reads.

### RNA-seq informatic analyses

Briefly, RNA-seq analysis was conducted through the following methods. Quality of FASTQ files from RNA sequencing were assessed using FastQC v.0.11.5 (Andrews, 2010). Adaptor sequences and low-quality reads were trimmed using Trimmomatic v.0.36 (Bolger et al., 2014). Next, alignment was performed using the STAR v.2.5.1a aligner (Dobin et al., 2013) to map reads to the mouse reference transcriptome, GRCm38.p6, to create a count matrix. We used DESeq2 (Love et al., 2014) to normalize the read count matrix, and to perform differential analysis. All statistical analyses and plots were generated using R v3.4.3. Pathview v.1.20.0 package was used for gene set enrichment analysis utilizing the KEGG database (Luo and Brouwer, 2013). KEGG pathways were classified manually using the BRITE functional hierarchies (Aoki-Kinoshita and Kanehisa, 2007) as an initial frame of reference, and final categorization was based on a consensus among three authors.

